# High-sensitivity c-reactive protein and total and saturated fat intake in adolescent students: a longitudinal study

**DOI:** 10.1101/349761

**Authors:** Camila Cândida de Lima Martins, Flávia Emília Leite de Lima Ferreira, Aléssio Tony Cavalcanti de Almeida

## Abstract

**Objective:** The present study aimed to assess the relationship between hs-CRP concentrations and total and saturated fat intake in adolescents after a year of follow-up.

**Methods:** A longitudinal study carried out in the years 2014 and 2015 evaluated 408 adolescents from the municipal and state public schools of João Pessoa, Paraíba, between 10 and 14 years of age, who participated in the Longitudinal Study on Sedentary Behavior, Physical Activity, Eating Habits and Adolescent Health (LONCAAFS). Data were obtained on sociodemographic data, anthropometric nutritional status, physical activity and hs-CRP concentration. The consumption of total and saturated fats was evaluated from the 24 hour recall.

**Results:** The associations between concentrations of hs-CRP and total and saturated fat intake were performed by linear regression considering panel data, individual fixed effect, balanced bank, stratified by sex and BMI. The mean values of the hs-CRP variable were significantly different between the analyzed years (p = 0.024). The percentage of total and saturated fat consumption is within the recommended level in both years, with no significant difference (p> 0.05). No statistically significant associations were found between hs-CRP and total fat consumption (β = −0.19p = 0.582) and saturated fat (β = 0.20, p = 0.282).

**Conclusion:** The study did not present significant evidence on the relationship between the concentrations of hs-CRP and the consumption of total and saturated fats, as one year of follow-up may not have promoted evident changes in the levels of hs-CRP in adolescents.

## Introduction

Inflammation is a natural body response to treatments or infections, aimed at promoting tissue regeneration [1]. Low-grade inflammation is associated with several non-communicable diseases, including obesity [2], diabetes mellitus, and metabolic syndrome [3]. Moreover, it is understood as an essential pathogenic mechanism at the onset and development of cardiovascular diseases (CVD) [4]. Despite the fact that this disease manifests during adulthood, the atherosclerotic process begins during childhood [5].

Particular attention has been given to inflammatory markers due to their capacity to predict the risk of CVD [6]. C-reactive protein (CRP) is an acute-phase inflammatory marker produced by the liver as a response to interleukin 6 (IL-6), which is stimulated in its turn by the tumor necrosis factor alpha (TNF-alpha) [7].

Aiming to detect future cardiovascular changes, high-sensitivity c-reactive protein (hs-CRP) has been explored as an inflammatory marker, as high concentrations of this marker were found to be associated with arterial changes in children [8] and adolescents [9–10], thus suggesting a possible role of low-grade inflammation in the onset of atherosclerosis [11]. As it detects lower serum CRP concentrations than those obtained by more traditional laboratory methods [12], the hs-CRP not only plays the role of marker, but also another role in the physiopathology of CVD [13].

The association between hs-CRP concentrations and fat intake has been identified in children, showing that both the total fat intake and percentage of energy from fat were positively associated with CRP concentrations and that the recurrent intake of foods considered to be “inflammatory” is directly associated with changes to hs-CRP concentrations in children [14].

A meta-analysis performed from intervention studies identified that the reduction in [15]. A meta-analysis performed with randomized trials suggests that a decrease in saturated fat intake promotes a reduction in the risk of development of CVD [16]. On the other hand, the intake of saturated fatty acids can increase inflammation and resistance to insulin, two determining factors associated with the development of atherosclerosis in adolescents with obesity [17].

There are gaps in the literature regarding the relationship between hs-CRP concentrations and the intake of macronutrients, especially fats in adolescents. The majority of publications are aimed at exploring the associations between hs-CRP and BMI, rather than directly through the intake of foods or specific macronutrients [18].

The present study aimed to assess the relationship between hs-CRP concentrations and total and saturated fat intake in adolescents living in a city in Northeastern Brazil.

## Materials and methods

### Population and sample

A longitudinal study was performed, based on data from the *Estudo Longitudinal sobre Comportamento Sedentário, Atividade Física, Hábitos Alimentares e Saúde de Adolescentes* (LONCAAFS - Longitudinal Study on Adolescents’ Sedentary Behavior, Physical Activity, Eating Habits and Health). This study began in 2014 and ended in 2017 was with adolescents of both sexes, aged between ten and 14 years, enrolled in the 6^th^ grade of public primary schools in the city of João Pessoa, state of Paraíba, Northeastern Brazil. It aimed to analyze the interrelationships among physical activity level, sedentary behavior, eating habits, quality of life and health indicators of adolescents, based on interviews, anthropometric measurements and biochemical tests. The exclusion criteria adopted for data analysis were as follows: adolescents out of the age group of interest for this study (<10 and >14 years), pregnant and/or lactating adolescents and with physical and mental disabilities that impeded participation in the study. Data from 2014 and 2015 were used in the present study.

The following parameters were taken into consideration to calculate sample size: 95% confidence interval, acceptable error of 4%, and design effect (deff) of 2. The minimum initial sample size was 1,130 adolescents, with an additional 40% to compensate for losses and refusals, resulting in 1,582 adolescents in the initial sample in 2014.

Single-stage cluster sampling was performed to obtain the sample from the LONCAAFS Study, where 28 schools were systematically selected (14 city and 14 state schools), distributed proportionately according to size (number of students enrolled in the 6^th^ grade) and geographical area of the city (North, South, East and West).

The present study used a representative sub-sample of the population, resulting in 17 schools which were submitted to post-stratification, following the technical recommendations in all areas. Thus, a group was formed, proportionately distributed by size (number of students enrolled in the 6^th^ grade) and geographical area of the city (North, South, East and West) (Figure 1).

**Figure 1:**
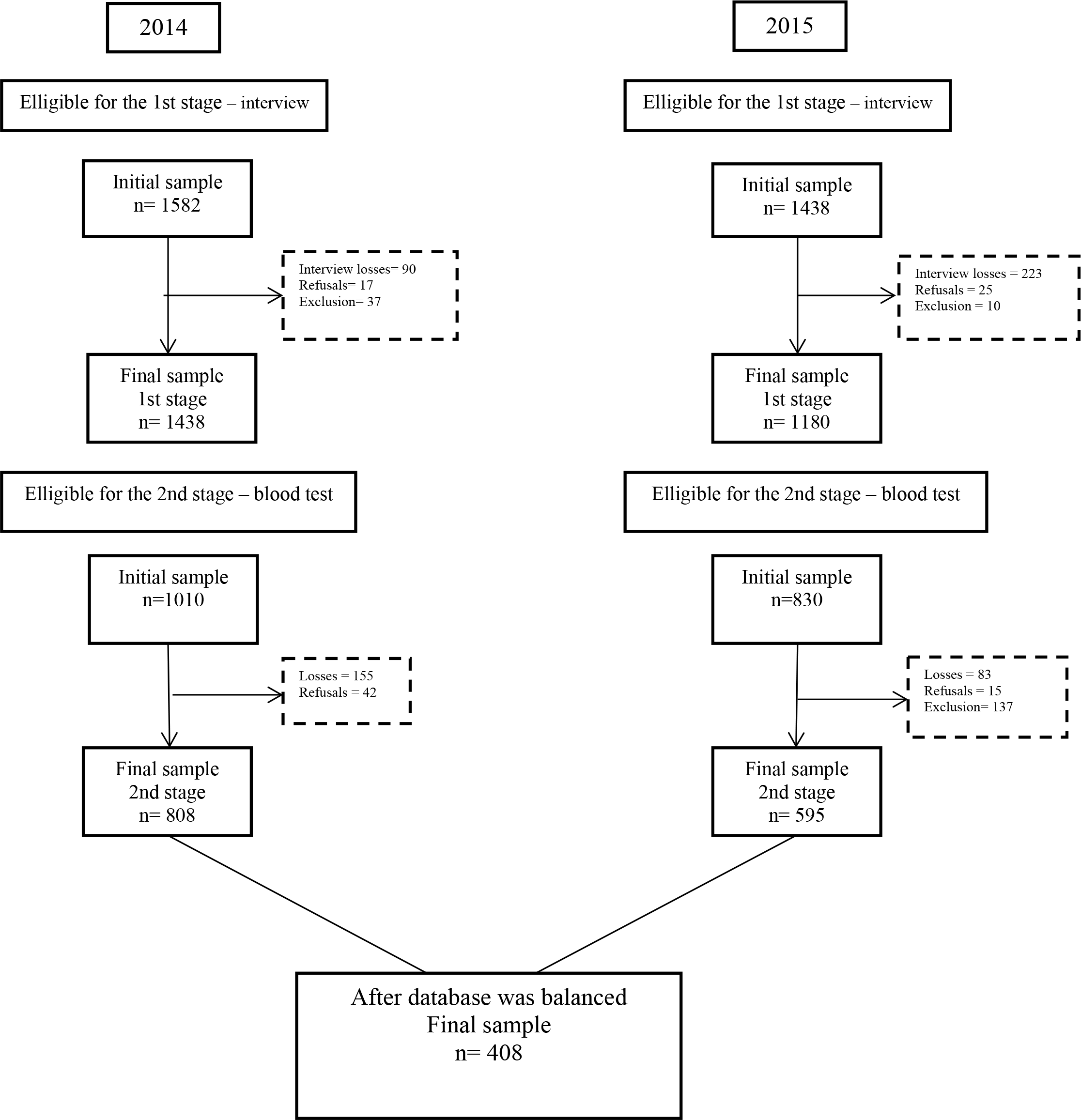
Flow chart of the 2014 and 2015 LONCAAFS study sample

### Ethical questions

The present research project was approved by the Human Research Ethics Committee of the Health Sciences Center of the Federal University of Paraíba. Aiming to collect data, parents or legal guardians of adolescents aged less than 18 years signed two informed consent forms to agree with the participation of adolescents in this study: one to be interviewed and the other to allow blood to be collected.

### Procedures

Data from the LONCAAFS Study from 2014 and 2015 were collected by the team comprised of nutritionists and physical educators and graduate and undergraduate Nutrition and Physical Education students from the UFPB.

Data collection was divided into two moments. The first moment involved the application of a questionnaire, performed through a face-to-face interview, when students were invited to provide information about socio-demographic variables, physical activity and food intake. Anthropometric measurements (weight and height) were taken to calculate BMI/age. The second moment, which occurred between one to two weeks after the first one, involved blood collection for hs-CRP analysis. Both stages were performed in the school itself. The sub-sample was exclusively comprised of adolescents who participated in both moments of data collection.

### Study Variables

The variables selected for the present study were as follows: socio-demographic factors, physical activity, weight, height, eating habits and hs-CRP measurement.

#### Sociodemographic variables

the socio-demographic variables analyzed were sex (male and female); age, ethnic group classified according to the *Instituto Brasileiro de Geografia e Estatística* (IBGE - Brazilian Institute of Geography and Statistics) [19] - white, mixed, black, Asian-descendant and indigenous –, maternal level of education, socioeconomic class according to the criteria recommended by the *Associação Brasileira de Empresas de Pesquisa* (ABEP - Brazilian Association of Market Research Companies) [20].

#### Physical activity

The questionnaire about physical activity, validated for the study population, was comprised of a list with 19 types of physical activity, with the possibility of including other unquestioned physical activities. Adolescents were encouraged to report the physical activities practiced by them in the previous week, informing the frequency in number of days per week duration of practice. The score was obtained by multiplying the number of minutes per day spent on physical activity by the number of days per week of each activity and subsequently adding the result of 19 activities, thus leading to a total score of physical activity per week. The classification of physical activity from the WHO Global Recommendations [21] considers that the practice of ≤ 300 minutes/week is regarded as physically inactive; and >300 minutes/week, physically active.

#### Body Mass Index

Aiming to diagnose the anthropometric nutritional status, the body mass index (BMI) was used, determined according to height and weight measurements and classified according to the criteria recommended by the World Health Organization (WHO), with categorization by age and sex [22]. Body mass (in kg) was observed using a Bioland^®^ digital scale with a capacity to measure between 2kg and 150kg and an accuracy of 100g. Height was measured with a Sanny^®^ portable stadiometer with a capacity to measure up to two meters and an accuracy of 1mm, following the standards described by Lohman, Roche and Martorell [23]. The procedures were performed in duplicate and a third measurement was taken in case the result of the second measurement was different from that of the first one. The values for the classification of anthropometric nutritional status [22] considered the information about Z-scores of BMI [(individual value − mean)/ standard deviation].

#### Food intake

Food intake was assessed with the use of the 24-hour dietary recall (24HR), applied once in the entire sample and with 30% replicated each year os the study. [24]. The Multiple Pass Method (MPM) was used to administer the 24HR [25]. The 24HR were applied by undergraduate and graduate Nutrition students from the Federal University of Paraíba (UFPB) and by volunteer nutritionists. Adolescents provided information about foods, ways to prepare, brand of manufactured foods and the amount consumed. When difficulties to measure the portions of certain foods were encountered, a photo album with images of standard cooking measures and varied food portions was used as reference, aiming to help to estimate individual intake and reduce memory bias [26]. For the analysis of the present study, only energy (kcal), total fat and saturated fat were assessed, characterizing a total fat intake ≥25% and ≤35% as adequate and <25% and > 35% as inadequate and a saturated fat intake <10% as adequate and ≥10% as inadequate, according to the Dietary Guidelines for Americans (2015-2020) [34]. Data on food intake were exported to an Excel spreadsheet, where their nutrients were shown.

#### hs-CRP analyzes

hs-CRP was determined in serum with the high-sensitivity ELISA method, using the Labtest’s Ultra Turbiquest Plus kit^®^. Analyses were performed with Labtest’s Labmax 240 Premium automated biochemical analyzer. Prior to each sequence of analysis, the equipment was calibrated with Labtest’s Calibra, aiming to assess the accuracy of estimates of hs-CRP and to observe whether they were within the parameters recommended by the equipment manufacturer. Analyses were performed in the Laboratory of Physical Training Applied to Development and Health of the Department of Physical Education at the Federal University of Paraíba. The classification of US-CR, categorized into unaltered (hs-CRP <1.0 mg/L) and altered (≥1.0mg/L), was performed according to the recommendations of the National Academy of Clinical Biochemistry [27].

### Data processing

The data collected were tabulated with the EpiData 3.1 software. Double-data entry was performed with subsequent automatic checking of consistency of responses using the “double data entry validation” tool to identify typing errors. These errors were identified and then corrected according to the values contained in questionnaires. Data on food intake, obtained with the 24HR, were tabulated and processed using the Virtual Nutri Plus^®^ software. In order to develop the food database used in the Virtual Nutri Plus^®^ software, in addition to those that had already been registered, other foods not consumed routinely or regionally were added to the food database of the software according to the Brazilian Food Composition Table [28] and the table for the assessment of food intake in cooking measures [29]. The correction of intra-individual variability of dietary recalls was performed with the Multiple Source Method (MSM). The MSM is a statistical method to estimate individuals’ usual dietary intake, including the sporadic intake of certain foods applying the 24HR two or more times. The method includes three stages: in the first stage, the probability of food intake is estimated per day for each individual. In the second stage, the usual food intake is estimated on the days of consumption. Finally, in the third stage, the usual food intake on all days is calculated by multiplying the probability of consumption by the usual amount of food intake on days of consumption [30].

### Statistical analysis

Descriptive statistics was performed to characterize the sample, showing the mean and standard deviation for continuous variables, frequency and percentage for categorical variables. The association between categorical variables and observed time (2014 and 2015) and the relationship between altered and unaltered hs-CRP and adequate and inadequate total fat intake were performed with the Chi-square test (χ^2^). Paired t-test was conducted in continuous variables to compare means between the observed years. The association between hs-CRP concentrations and fat intake in adolescents was assessed with multiple linear regression, including panel data with an individual fixed effect and balanced database, controlled for confounding factors and stratified by sex and BMI. The hs-CRP corresponded to the dependent variable in the study, whereas total fat intake and saturated fat intake were the independent variables. BMI/age, socioeconomic class, physical activity and energy expenditure in kcal were the control variables. All variables were transformed into a natural logarithm, as they did not show a normal distribution. Sex and age were considered to be fixed variables, as they did not vary between the years assessed. All analyses took into consideration a 95% significance level. In order to obtain the results, the database was previously balanced, aiming to observe the existence of temporal data on the same individuals in 2014 and 2015. The use of the fixed effect is convenient for analyses with variables that change with time, as it enables one to study the causes of changes in the same individuals [31]. All analyses were performed with the STATA 13.0 software. After the database for 2014 and 2015 was balanced with an eligible sample for the second stage, the final sample was comprised of 408 individuals who showed all the variables analyzed in both years.

## Results

The variables that characterize the sample are shown in Table 1. Women comprised 56% of the population, between 10 and 11 years of age, presenting mostly a middle and low income class. Regarding the BMI / age of adolescents around 65% of the sample is found to be without excess weight.

**Table 1.**
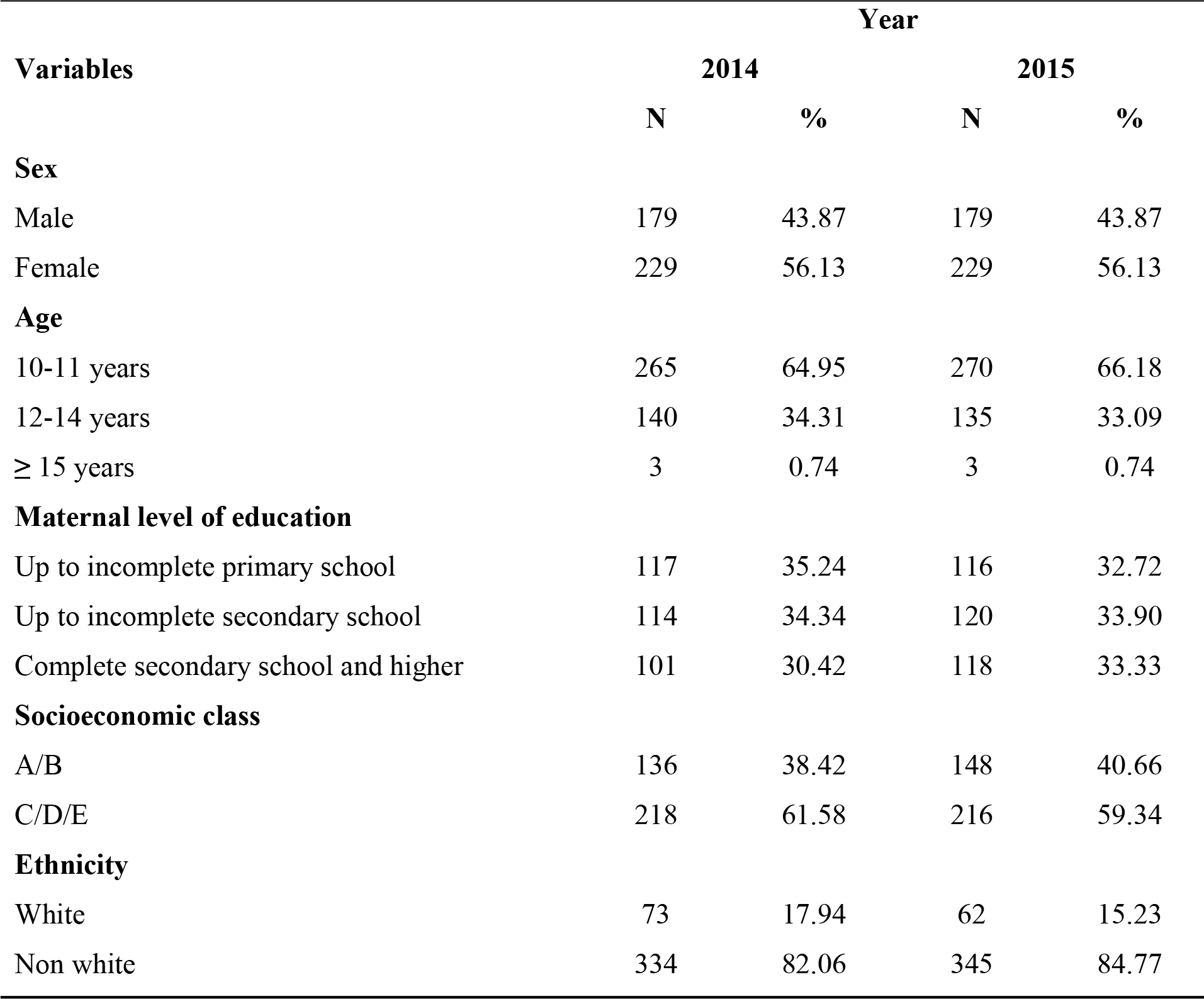
Characterization of socio-demographic variables and BMI of adolescents, 2014 and 2015.

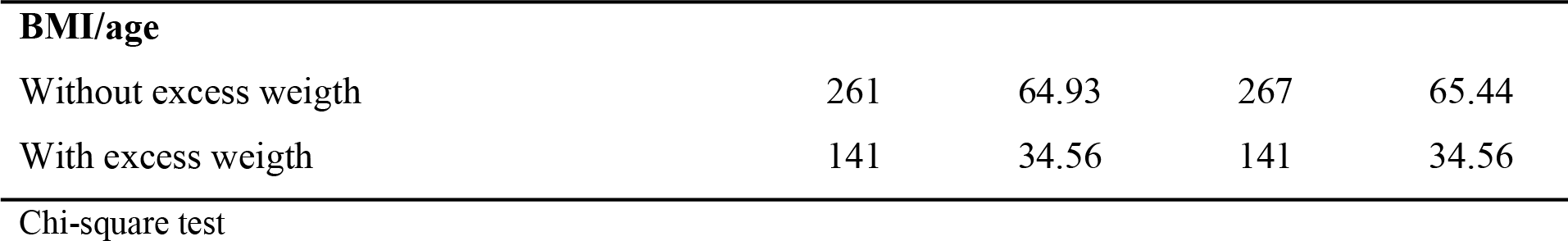

Table 2 shows the data on mean and standard deviation for control variables, according to the time observed. The hs-CRP showed a statistically significant difference between the means of the years analyzed (p<0.05). There was no significant difference between the years analyzed for total fat consumption, as well as for saturated fat (p> 0.05). Analyzing the average percentages of consumption, they were found to be within the nutritional recommendations for the evaluated public.

**Table 2.** Mean and standard deviation of variables according to the observed time in adolescents, 2014 and 2015

| Variable | Time | Mean | SE | 95%CI | P |
| --- | --- | --- | --- | --- | --- |
| **BMI (kg/m<sup>2</sup>)** |  |  |  |  |  |
|  | 2014 | 19.78 | 0.21 | 19.36 - 20.20 | 0.489 |
|  | 2015 | 20.57 | 0.23 | 20.12 - 21.05 |  |
| **Energy (kcal)** |  |  |  |  |  |
|  | 2014 | 2322.10 | 65.93 | 2193.56 - 2452.43 | 0.483 |
|  | 2015 | 2172.99 | 35.03 | 2104.21 - 2241.76 |  |
| **Total fat (%)** |  |  |  |  |  |
|  | 2014 | 29.33 | 0.57 | 28.21 - 30.45 | 0.244 |
|  | 2015 | 29.28 | 0.28 | 28.72 - 29.83 |  |
| **Saturated fat (%)** |  |  |  |  |  |
|  | 2014 | 7.57 | 0.27 | 7.05 - 8.09 | 0.494 |
|  | 2015 | 7.73 | 0.20 | 7.49 - 7.96 |  |
| **hs-CRP\* (mg/L)** |  |  |  |  |  |
|  | 2014 | 0.98 | 0.10 | 0.79 - 1.18 | 0.024 |
|  | 2015 | 1.47 | 0.12 | 1.23 - 1.70 |  |
| **Physical** |  |  |  |  |  |

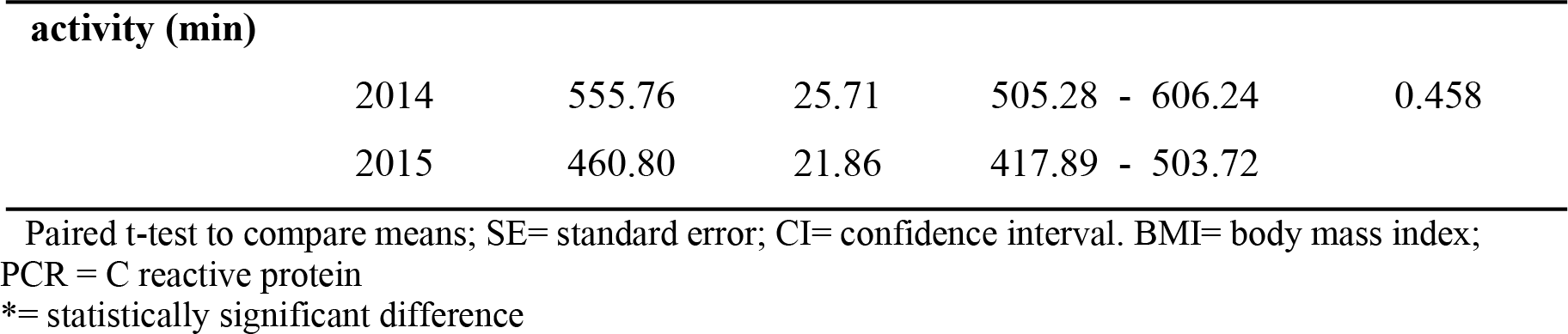

As shown in Table 3, the relationship between the consumption of adequate and inadequate saturated and total fat with altered and unchanged concentrations of hs-CRP is observed, and we identified that around 70% of the 2014 sample and 80% of the sample of 2015 showed adequate intake of both total fat and saturated fat and adequate hs-CRP levels, but the relationship between the variables did not present statistical significance (p> 0.05).

**Table 3.** Relationship between hs-CRP and saturated and total fat consumed by adolescents, 2014 and 2015.

|  |  | hs-CRP |  |  |  |  |  |
| --- | --- | --- | --- | --- | --- | --- | --- |
|  |  | Unaltered |  | Altered |  | Total | % |
| 2014 |  | n | % | n | % | n |  |
| Total fat (%) | Adequate | 231 | 78.31 | 79 | 79.80 | 310 | 78.68 |
|  | Inadequate | 64 | 21.69 | 20 | 20.20 | 83 | 21.32 |
| p |  | 0.754 |  |  |  |  |  |
| Saturated fat (%) | Adequate | 217 | 73.56 | 73 | 73.74 | 290 | 73.60 |
|  | Inadequate | 78 | 26.44 | 26 | 26.26 | 104 | 26.40 |
| p |  | 0.972 |  |  |  |  |  |
| 2015 |  |  |  |  |  |  |  |
| Total fat (%) | Adequate | 217 | 87.50 | 141 | 88.68 | 358 | 87.96 |
|  | Inadequate | 31 | 12.50 | 18 | 11.32 | 49 | 12.04 |
| p |  | 0.721 |  |  |  |  |  |
| Saturated fat (%) | Adequate | 208 | 83.87 | 129 | 81.13 | 337 | 82.80 |
|  | Inadequate | 40 | 16.13 | 30 | 18.87 | 70 | 17.20 |
| p |  | 0.475 |  |  |  |  |  |

Chi-square test – without statistical significance ( $p > 0.05$ )

The crude and adjusted analyses of linear regression were performed to analyze the association between hs-CRP concentrations and total and saturated fat intake. There were no statistically significant associations between variables (p>0.05), as observed in Table 4.

**Table 4.**
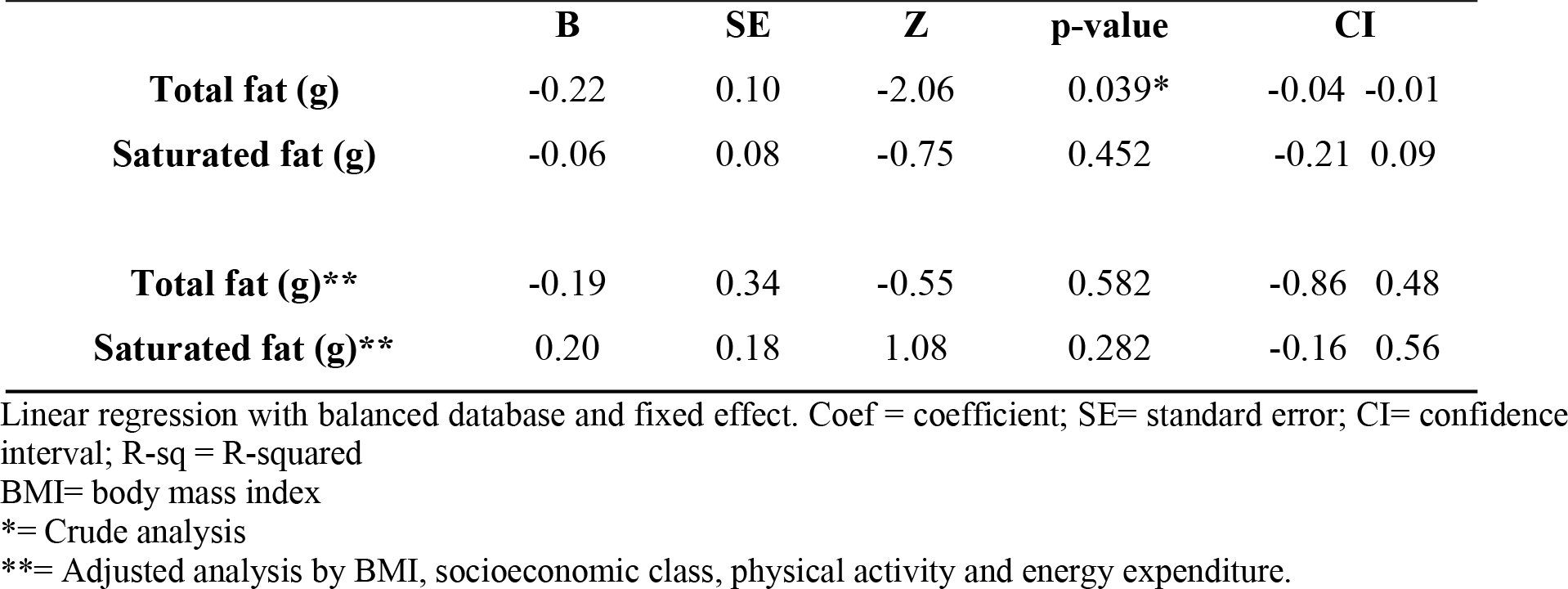
Analysis of crude and adjusted linear regression between hs-CRP concentrations and total and saturated fats consumed by adolescents, 2014 and 2015.

|  | **B** | **SE** | **Z** | **p-value** | **CI** |
| --- | --- | --- | --- | --- | --- |
| **Total fat (g)** | -0.22 | 0.10 | -2.06 | 0.039* | -0.04 -0.01 |
| **Saturated fat (g)** | -0.06 | 0.08 | -0.75 | 0.452 | -0.21 0.09 |
| **Total fat (g)\*\*** | -0.19 | 0.34 | -0.55 | 0.582 | -0.86 0.48 |
| **Saturated fat (g)\*\*** | 0.20 | 0.18 | 1.08 | 0.282 | -0.16 0.56 |

Linear regression with balanced database and fixed effect. Coef = coefficient; SE= standard error; CI= confidence interval; R-sq = R-squared
BMI= body mass index
\*= Crude analysis
\*\*= Adjusted analysis by BMI, socioeconomic class, physical activity and energy expenditure.

The Table 5 presents data from the adjusted linear regression analysis performed to evaluate the concentrations of hs-CRP with total and saturated fat consumption stratified by sex and BMI / age. No statistically significant difference was observed (p>0,05).

**Table 5.**
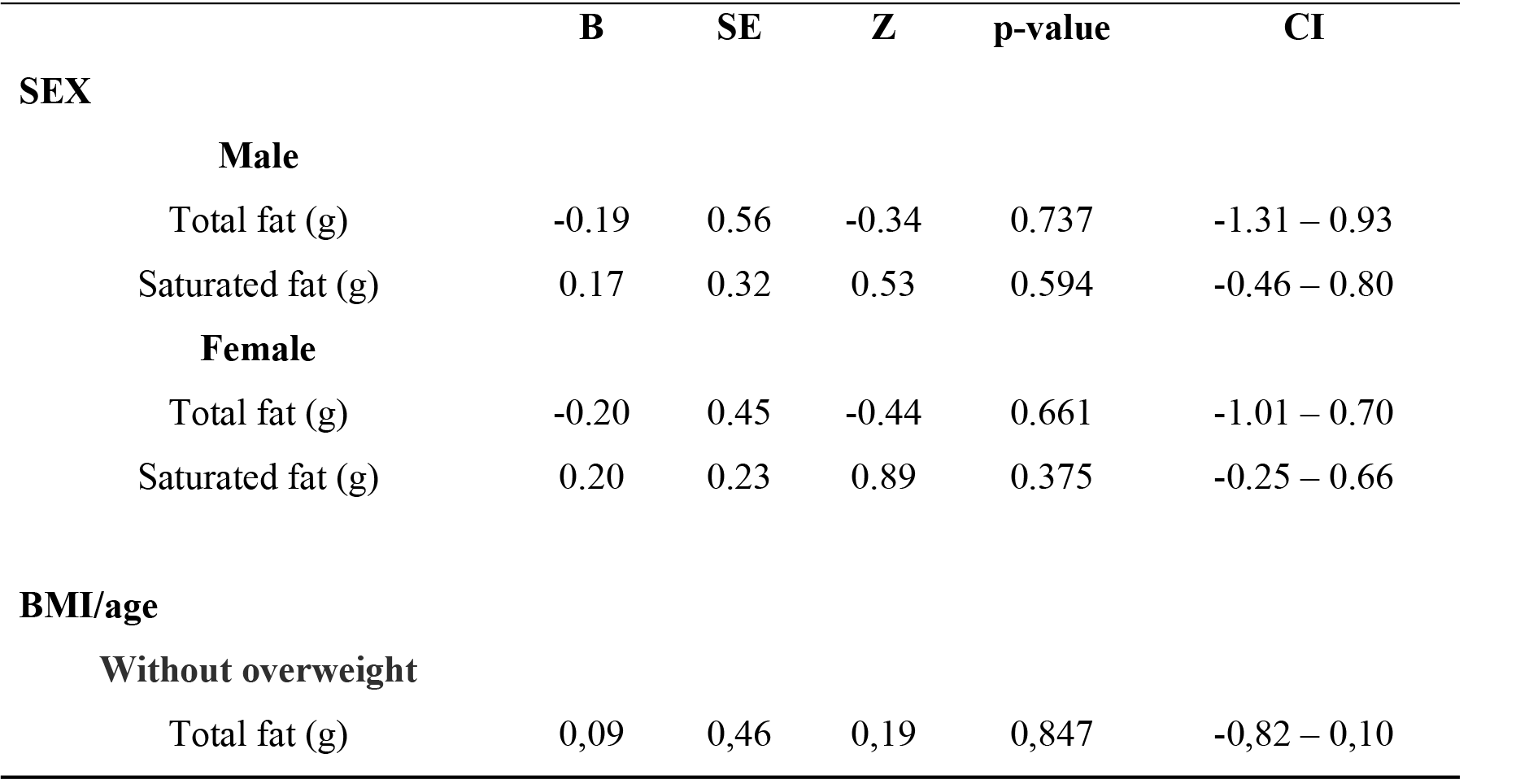
Adjusted linear regression analysis between hs-CRP concentrations and total and saturated fat consumed by adolescents 2014 and 2015, stratified by sex and nutritional status.

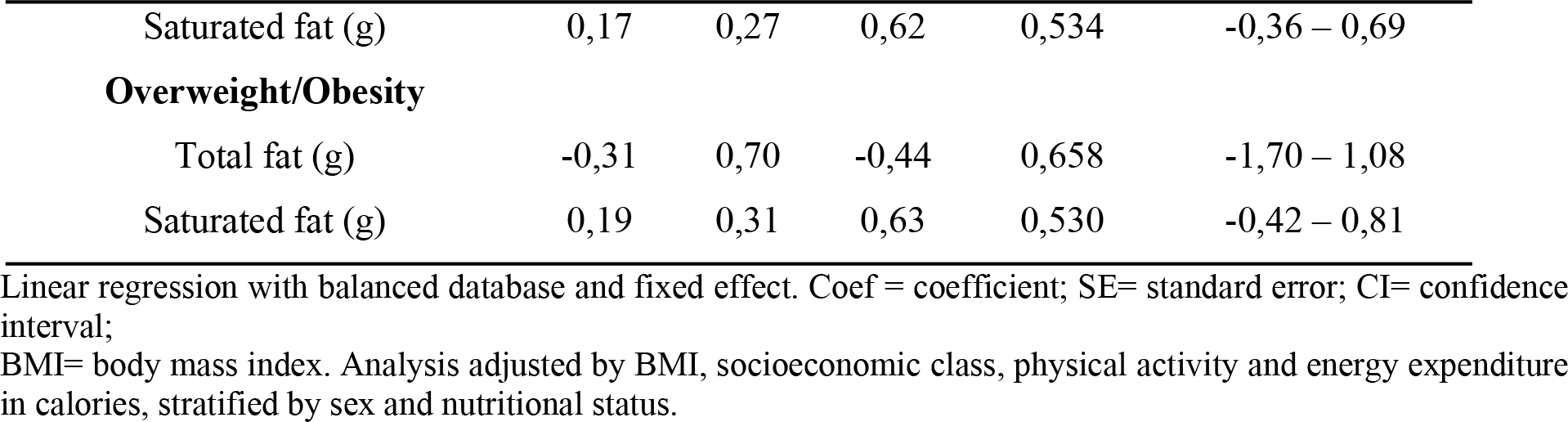

## Discussion

According to the sample characteristics, the majority of participants were female adolescents aged between 11 and 12 years from socioeconomic classes C/D/E, thus indicating a low- and medium-income population. The percentage of total and saturated fat intake did not show significant differences. However, when the mean values from 2014 and 2015 are observed, total fat intake was found to meet the recommendations of the Brazilian Cardiology Society [32], Ministry of Health [33] and *Dietary Guidelines for Americans* (2015-2020) [34].

According to the *Pesquisa de Orçamento Familiar* 2008-2009 (POF - Family Budget Survey), regardless of one’s income level, total fat intake was found to be adequate as observed in our study [35]. The same can be observed for saturated fat intake and, although not showing significant differences between the years analyzed, the percentage consumed by adolescents was lower than the recommendations, totaling <10% of the total energy intake among adolescents aged between nine and 13 years [32–34].

These data point to values below those found by the *Estudo dos Riscos Cardiovasculares em Adolescentes* (ERICA - Study on Cardiovascular Risks in Adolescents) Study, which showed a mean total fat intake of 31.0% for girls and 30.0% for boys and a total saturated fat intake of 11.0% for both sexes, thus being higher than the recommendations for such adolescent population. The same was seen when compared to the *Inquérito Nacional de Alimentação* 2008-2009 (INA - National Dietary Survey), which although not presented results superior to the recommended for adolescents, as in ERICA, presented a consumption of total fats of 27% and of saturated fat 10% [36].

The mean hs-CRP concentrations in adolescents in the present study in the city of João Pessoa changed from low-to medium-risk with the change of time. In the study developed by Giannini et al. [37] with adolescents from six capital cities in Brazil using data from the ERICA Study it can be observed that the mean of CRP for the study in the city of João Pessoa considered altered ranged between 0.20 and 3, 20 according to the percentiles division (50 to 90), and that this increase was more evident in adolescents with excess weight, ranging from low to high risk, higher than that found in the present study.

When observing the adequate or inadequate consumption of total fats and saturated with unchanged and altered concentrations of hs-CRP, it is identified that both consumption was found to be adequate for total fats and saturated in both 2014 and 2015. Adequate fat consumption observed in the two-year follow-up may be linked not to a balanced diet but to a monotonous diet low in complex carbohydrates and rich in simple sugars and fats, which is often associated with obesity and other diseases non-communicable chronic diseases, mainly seen in the low-income population [38].

Data on the relationship between total fat intake and inflammatory markers such as the hs-CRP in adolescents are still scarce [9,39–40]. In a study performed with adolescents aged more than 13 years, Lazarou et al.[41] did not find a statistically significant association between quality of diet and inflammation through hs-CRP concentrations. However, this study found a strong association between obesity (increase in BMI/age) and higher hs-CRP concentrations.

Lee, Gurka and Deboer [42] investigated the relationship between metabolic syndrome and cardiovascular risks in a population aged from 12 to 20 years, associating hs-CRP with the intake of total fats, carbohydrates, proteins and physical activity. They observed that the metabolic syndrome is directly associated with cardiovascular risk, although a relationship between the hs-CRP concentrations and fat intake analyzed in this study was not found.

The consumption of different types of fat contributed to the increase in hs-CRP concentrations in European children [40]. In Brazil, a study performed with children and adolescents aged from five to 13 years revealed an association between the increase in the intake of high-fat processed foods and higher hs-CRP concentrations. Nonetheless, it is not clear whether the relationship was more significant in children or in adolescents [14].

The majority of the population studied was within the normal hs-CRP concentration parameters. It is believed that the lack of association between this concentration and total fat intake may have been influenced by other types of fat that were consumed but not analyzed, such as monounsaturated and polyunsaturated fats.

Adherence to a diet based on anti-inflammatory components, such as the Mediterranean diet, showed a decrease in the risk of cardiovascular and cerebrovascular diseases [43]. A randomized clinical trial with female adolescents with metabolic syndrome assessed the influence of the Dietary Approaches to Stop Hypertension (DASH) diet on inflammatory marker levels for six weeks. It concluded that higher consumption of vegetables, fruits and whole grains, low-fat and fat-free dairy products, fish, poultry, beans, nuts and vegetable oils and lower consumption of high-saturated fat foods, sweetened beverages and candies contribute to the reduction in serum hs-CRP concentrations [44].

Studies point to circulating CRP concentrations are associated with higher BMI [45]. Halder et al. [46] and Bochud et al. [47] reported that CRP values rose with the increase in BMI and that obese individuals show a high CRP prevalence more frequently.

Although a slight increase in BMI occurred between 2014 and 2015, it was not associated with altered CRP concentrations in the present study. A comparison made between overweight and eutrophic children found that those who were overweight showed higher CRP concentrations and a seven times higher risk of developing metabolic syndrome in adulthood if they become obese adults [48]. Roh et al. [49] pointed out that, when adolescents became obese, they showed inflammatory characteristics that were similar to obese adults.

However, the lack of association in this study does not dismiss the importance of continuing the analysis of such association, as the hs-CRP is associated with cardiovascular risk factors [50] and the fat intake in this population has increased. However, the existing results originate from cross-sectional studies, as indicated by a study that assessed two groups of children and adolescents aged between six and 12 years and found an increase in the consumption of high-total and saturated fat foods [51], as observed by Leal et al.[52] and Silva et al.[53]

The present study had some positive points and limitations. Until now, this has been the first study in Brazil that associated hs-CRP concentrations and total and saturated fat intake in adolescents. It should be emphasized that only two years of follow-up were assessed and that future analyses will be able to evaluate four years of study, aiming to verify this association throughout the entire first stage of adolescence. Panel data were analyzed in a balanced way and, although involving a reduction in the final sample size, the balanced analysis is more accurate. Moreover, the assessment of unsaturated fats is also necessary, as they have an anti-inflammatory potential that can interfere with hs-CRP results.

During the application of the 24HR, memory bias can lead to under- or overestimation in the reports of intake of certain foods, hindering the quantification of nutrients of interest in the study and energy consumption in calories. Likewise, not reporting the intake of fats used in preparations, as many adolescents had difficulty describing how foods were prepared, may have led to an underestimation of consumption. However, the 24HR is still the pattern to assess food intake in population studies. Another point that should be emphasized as a limitation was the lack of assessment of the maturing stage of adolescents, which could be associated with the inflammatory profile [54].

## Conclusion

There was a significant increase in hs-CRP concentration from 2014 to 2015, although total and saturated fat intake met the recommendations for the study population.

The present study did not show significant evidence of the relationship between a possible change in hs-CRP concentrations and total and saturated fat intake in the sample studied, regardless of sex and BMI.

The present findings provides guidance for future studies, suggesting the need for follow-up of adolescents during a longer period of time. Furthermore, it emphasizes the need to assess food intake not only through other nutrients such as carbohydrates, but also through foods and food groups.

## Acknowledgements

Authors would like to thank the entire LONCAAFS team for their commitment during data collection; the adolescents who participated in this study for their availability; and the funding institutions, CNPq and FAPESq.

